# gffutilsAI: an AI-agent for interactive genomic feature exploration in GFF files

**DOI:** 10.64898/2025.12.02.690645

**Authors:** Virginia Gonzalez, Tristan Yang, Sebastian Bassi

## Abstract

The General Feature Format (GFF) is widely used to represent genomic annotations, but its hierarchical, multi-attribute structure makes manual querying and analysis challenging. Existing libraries such as gffutils provide programmatic interfaces, yet they require coding proficiency. gffutilsAI is a novel AI-powered command-line agent that enables researchers to perform interactive, natural-language-driven exploration of GFF files. Built on top of the gffutils library and the Strands AI agent framework, gffutilsAI integrates local and cloud-based large language models (LLMs) such as Llama 3.1, GPT-5, and Claude 3.5 to translate human queries into executable actions. The tool supports coordinate-based queries, attribute and GO searches, hierarchical traversal, statistical summaries, and CSV export, offering a new paradigm for accessible conversational genomics.

## 1 Introduction

Artificial Intelligence (AI) applications in bioinformatics emerged in the mid-1990s, initially focusing on DNA sequencing, data management, and data mining using techniques such as genetic algorithms, hidden Markov models, and neural networks [1]. This landscape has been significantly transformed by the recent advent of transformers and LLM. Current AI applications have expanded to include DNA/RNA/protein sequence analysis and functional prediction [2], protein and RNA structure prediction [3, 4], transcriptomics data analysis [5], virtual screening and drug-target prediction [6, 7], multi-omics data analysis [8], and biomedical literature mining [9]. Virtually every bioinformatics subfield now features AI-driven approaches that perform on par with or exceed traditional methods [10].

Despite this broad integration, a notable gap remains in the application of AI for the orchestration and intuitive control of these specialized tools. Many powerful bioinformatics utilities require significant technical expertise, limiting their accessibility. One such utility is gffutils [11], which is a tool that provides programmatic access to gff files from Python. Here we introduce gffutilsAI, a proof-of-concept AI tool that allows researchers to perform interactive, natural-language-driven exploration of GFF files. GffutilsAI demonstrates how LLMs can be leveraged to create a natural language interface for complex bioinformatics libraries, thereby simplifying workflows and lowering the barrier to entry for researchers.

GFF and its modern version GFF3 are standards for representing genes, transcripts, and regulatory elements. While the gffutils library provides Python APIs for indexing and analyzing GFF data, it imposes a learning curve on users unfamiliar with scripting. A simpler, interactive tool that allows describing analyses in plain language rather than code, is an alternative way to use this library.

In recent years, LLMs have demonstrated strong natural language understanding and reasoning ability. Coupling LLMs with structured bioinformatics tools opens new pathways for adaptive, query-driven research. We developed gffutilsAI, an AI agent that bridges human questions and structured genomic analysis, offering a hybrid workflow between scripting and conversation.

## 2 Availability

- Project name: gffutilsAI (install with `pip install gffutilsai`)
- Code repository: https://github.com/ToyokoLabs/gffutilsAI (DOI: 10.5281/zenodo.17796146)
- Operating systems: Linux, macOS, Windows
- Programming language: Python (3.12+)
- License: GPL-3.0

## 3 Implementation

### 3.1 Architecture

gffutilsAI extends gffutils Python library with an AI agent layer powered by Strands[12], an open-source framework for integrating LLM-based tools. See Figure 1 for a schematic of how the agent uses the model to select and execute a tool, passing the results back to the model to generate the final output. This integration bridges the gap between conversational English and structured genomic data, eliminating the need for users to write custom scripts for routine analysis. It includes more than 18 specialized tools available to the AI agent, which can be classified as:

**Figure 1.**
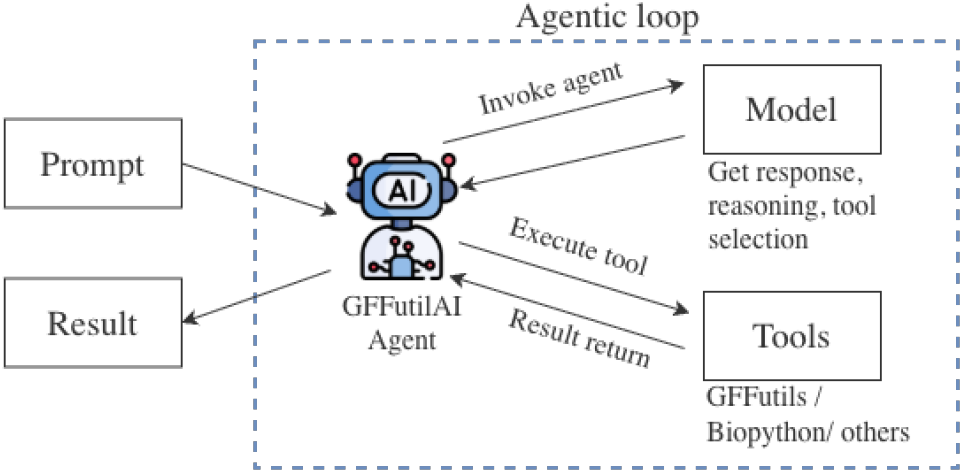
GFFUtilsAI Agent Workflow: The agent uses the model to select a tool, executes the tool, retrieves the output, and feeds it back to the model to generate the result.

- Coordinate-based: retrieve features overlapping specific genomic coordinates
- Structural: navigate hierarchical relationships such as exons, CDS, and genes
- Statistical: compute counts, length distributions, and feature densities
- Attribute-based: search by attribute values or patterns (e.g., Name, Note, GO_function)
- Positional: find intergenic regions, strand biases, and chromosomal organization
- Export & Reporting: produce CSV tables and text summaries for downstream analysis

Each tool is defined in gff_tools.py and uses the @tool decorator for exposure to the AI layer. The main.py script manages runtime parameters such as model selection, temperature, and token limits. The system supports inference through major cloud providers (OpenAI, Anthropic, Google) as well as local execution via Ollama. Theoretically, any Ollama-hosted model with tool support is compatible; however, our testing revealed that some models advertised as ‘tool-capable’ failed to perform reliably in practice (see Table 1 for a list of verified compatible models). Configuration is handled via command-line arguments or environment variables, and the system includes a batch mode for processing multiple questions non-interactively.

**Table 1:** Ollama compatible models that support tools and are known to work with gffutilsAI.

| **Model Name** | **Observation** |
|---|---|
| glm-4.6:cloud | Functions as expected |
| gpt-oss:20b-cloud | Functions as expected |
| gpt-oss (20b and 120b) | Functions as expected |
| llama3 (All variants) | Subpar results; even llama3-groq-tool-use yields low-quality output. |
| qwen3:14b | Successfully calls tools but fails to accurately interpret most user queries. |

## 4 Sample usage

Here are some sample queries to showcase different kind of questions about a gff file that the agent is capable to answer.

### 4.1 Coordinate-based Queries

This query requests a list of genes located in the final 8000 nucleotides of a specific chromosome. To address this, the LLM requires the chromosome’s length to determine the target coordinate range. After establishing the coordinates, it can search for the relevant genes. See how the agent uses three tools to complete this work:

~~~
List all genes available in the last 8000 nucleotides of the chromosome CM116928 . 1 in / content / GCA_050947715 . 1 _ASM5094771v1_genomic . gff .
------------------------------
I ‘ ll help you find the genes in the last 8000 nucleotides of the chromosome CM116928 . 1 . To do this , I ‘ ll first need to get the chromosome ‘s length and then use the get _ features _ in _ regionfunction to retrieve the genes .
Let ‘s break this down in to steps :
1. First , I ‘ll check the chromosome information :
Tool # 1 : get _ chromosomes_ inf o
2. Now, I ‘ ll get the chromosome summary to confirm the length :
Tool # 2 : get_chromosome_summary
The chromosome CM116928 . 1 has a length of 1 , 486 , 871 nucleotides . I ‘ll retrieve the genes in the last 8000 nucleotides:
Tool # 3 : get_features_in_region
I found 4 genes in the last 8000 nucleotides of chromosome CM116928 . 1 :
1. Gene ACO0OE_002496
  -Start: 1 , 478 , 846
  -End : 1 , 479 , 736
  -Strand: +
  -Length : 890 nucleotides
2. Gene ACO0OE_002497
   -Start: 1 , 479 , 918
   -End : 1 , 481 , 792
   -Strand: +
   -Length : 1 , 874nucleotides
3. Gene ACO0OE_002498
   -Start: 1 , 482 , 050
   -End : 1 , 482 , 919
   -Strand: +
   -Length : 869 nucleotides
4. Gene ACO0OE_002499
   -Start : 1 , 485 , 924
   -End : 1 , 486 , 115
   -Strand : +
   -Length : 191 nucleotides
All of these genes are protein  - coding genes ( gene _ biotype : protein _ coding )
and a relocatedon the positive strand ( + ) of the chromosome.
------------------------------
~~~

### 4.2 Statistical Analysis

This query demonstrates the agent’s ability to transition from simple data retrieval to quantitative profiling. The agent performs a multi-layered statistical assessment:

Comparative Sorting: It evaluates the chromosomal dataset to identify and isolate the “largest” entity (chromosome CM116930.1) based on sequence length.

Metric Derivation: Beyond simple counting, it computes derived statistics by normalizing raw counts into standard biological units, calculating Gene Density (470.85 genes/Mb) to allow for cross-genome comparisons.

Spatial Density Analysis: The agent utilizes a window-based approach (100 kb bins) to map the distribution landscape. It identifies the specific range of variation (430 to 590 genes/Mb) to empirically confirm that the genomic architecture is “relatively uniform.”

Notably, most LLMs in standalone mode fail to resolve this query due to the prohibitive size of GFF files and the extensive computation required to normalize density metrics across millions of base pairs.

~~~
What is the gene count and distribution pattern for the largest chromosome ?
--------------------------------------------------
I ‘ l l help you analyze the gene count and distribution for the largest chromosome in the GCA_050947715 . 1 _ASM5094771v1_genomic . gff file . I ‘ll break this down in to several steps :
1 . First , I ‘ll get the chromosomes to identify the largest one
2 . Then , I ‘ll get chromosome summary to understand its features
3 . Finally , I ‘ll analyze the gene density to show the distribution pattern
Let ‘sbegin :
1. Get Chromosome Information :
Tool # 1 : get _ chromosomes_info
2. Get Chromosome Summary to find the largest chromosome and its gene details :
Tool # 2 : get_chromosome_summary
3. Now, let ‘s get the gene density distribution for the larges t chromosome ( CM116930 . 1 ) :
Tool # 3 : get _ feature _ density
Analysis Summary for the Largest Chromosome ( CM116930 . 1 ) :
1 . Chromosome Length : 1 , 922 , 076 base pairs
2 . Total Gene Count : 905 genes
3 . Gene Density : 470 . 85 genes / Mb ( megabase )
Gene Distribution Pattern (in 100 kb windows ) :
 -Lowest gene density : 430 genes / Mb ( in windows 1 -100 kb and 300 -400 kb )
 -Highest gene density : 590 genes / Mb ( in the 1700 -1800 kb window )
 -Most windows have a gene density between 430 -540 genes / Mb
 -Slight variations in gene distribution across the chromosome
 -The last window ( 1900 - 1922 kb ) has a lower gene density due to its smaller size
Observations:
 -The gene distribution is relatively uniform across the chromosome
 -There are minor fluctuations in gene density , which is typical in genomic landscapes
 -The chromosome shows a consistent gene distribution pattern with no extreme clustering or depletion of genes
------------------------------
~~~

### 4.3 Other usage examples

Beyond the detailed workflows presented above, gffutilsAI handles a wide range of genomic queries. The following subsections highlight these diverse capabilities. For the sake of brevity, we omit the raw code and output listings here; readers can refer to the Supplementary Information at https://github.com/ToyokoLabs/gffutilsAI/blob/main/paper/Supplementary%20Information.pdf for the full interaction data corresponding to these examples.

#### 4.3.1 Relationship and Hierarchy Analysis

The agent has the capacity to navigate genomic topologies. It can reconstructs the hierarchical structure of the target locus, mapping the gene ACO0OE_002170 to its parent chromosome (CM116928.1) while identifying downstream child entities (mRNA, CDS, and Exons). In another query, it further tests this relational understanding; the agent correctly isolates a specific node in the hierarchy (the mRNA transcript) and quantifies its sub-components (CDS regions), accurately interpreting the one-to-many relationship between the transcript and its coding sequences.

#### 4.3.2 Retrieving Species Information

The agent correctly identified the organism as *Hanseniaspora uvarum*. This highlights an architectural adaptation addressing the lack of metadata extraction in `gffutils`: the system automatically invoked `get_organism_info` to query the NCBI Entrez database [13] via Biopython [14]. Standalone LLMs consistently failed this task, resulting in hallucinations of incorrect species.

#### 4.3.3 Attribute-based

The agent has an adaptive search capabilities when dealing with genomic metadata. When a strict, ontology-based query (search_genes_by_go_function_attribute tool) fails to yield results, the agent autonomously pivots to a broader attribute-based search. By querying the ‘product’ description field directly, it successfully identifies relevant proteins that lack specific GO annotations but contain the target keyword in their textual metadata.

#### 4.3.4 Positional

The agent is capable of defining intergenic regions of arbitrary size on any chromosome via the `get_intergenic_regions` tool. For instance, when asked to find regions exceeding 2500 bp on chromosome CM116926.1, it retrieved the top 10 entries and displayed their lengths, positions, and flanking genes in an easy-to-read format.

## 5 Comparative Analysis of Tools

To validate our approach, we performed a set of tests using the biological questions mentioned previously. To establish a performance baseline, we first evaluated a diverse set of 13 LLMs in a zero-shot configuration. Second, we tested different LLMs as the backend for gffutilsAI to observe how model selection affects agent performance.

### 5.1 Baseline Performance of Standalone Models

To evaluate how each LLM handles queries related to GFF files, we administered a six-question test based on a specific GFF dataset to 13 LLMs. The method for ingesting the file varied by model; some platforms supported direct file uploads, while others required command-line insertion due to interface constraints. The complete list of questions is detailed in Table 2, and all model responses are archived in the project’s software repository within the paper subdirectory.

**Table 2:** List of questions used to evaluate LLM performance on GFF file analysis.

| ID | Question Prompt |
|---|---|
| Q1 | I want a list of all attributes in GCA_050947715.1_ASM5094771v1_genomic.gff |
| Q2 | Tell me which species it is |
| Q3 | What is the geographical origin of this genome? |
| Q4 | Give me the number of chromosomes |
| Q5 | Give me the number of genes on each chromosome |
| Q6 | List all vacuolar proteins and their protein_id |

The performance in this control group was highly stratified (Figure 2). While the proprietary state-of-the-art model (ChatGPT5 Pro) achieved a perfect score (6*/*6), the majority of other models struggled significantly with the test set. Notably, high-parameter open-weight models such as Llama 3.3 70b, GLM 4.6, and gpt-oss:120b failed to answer a single question correctly (0*/*6). Mid-sized proprietary models like Haiku 4.5 and Gemini 2.5 Flash similarly underperformed, scoring only 1*/*6. This baseline demonstrates that without architectural scaffolding, standalone models face two distinct yet compounding bottlenecks. First, many lack the intrinsic reasoning chain required to navigate the complexity of these specific queries. Second, and perhaps more prohibitive, is the structural limitation of the context window. For several models, the raw genomic input (a GFF file) exceeded the available token limit, effectively preventing the model from ingesting the necessary data to formulate a response.

**Figure 2.**
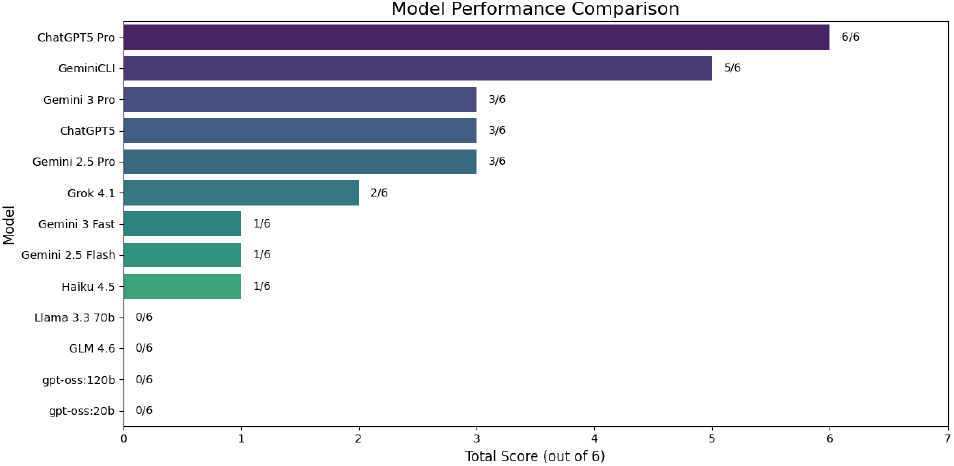
Baseline performance of standalone LLMs on the GFF analysis benchmark. The bar chart displays the total score (out of 6) for each of the 13 models tested in a zero-shot configuration.

### 5.2 How the LLM contributes to the GFFUtilsAI Agent’s performance

To evaluate the efficacy of the GFFUtilsAI agent across different architectures, we utilized the system’s batch processing mode to execute the benchmark question set while systematically varying the underlying LLM. A subset of the original models was selected for this analysis, as not all previously tested LLMs possess the requisite capabilities for agentic tool use. All experimental data are archived in the project repository, accompanied by a Jupyter notebook (`gffutilsaitest.ipynb`) to facilitate reproduction. We acknowledge that the inherent stochasticity of LLMs may introduce slight variance between runs; therefore, in instances of intermittent failure, the optimal successful response was recorded. Qualitatively, we observed that reproducibility within the agentic workflow was significantly more consistent than in the standalone mode (data not shown).

Figure 3 illustrates the contribution of the underlying LLM to the final agentic performance. While the agentic scaffolding enabled all tested models to achieve passing grades ( *≥* 4*/*6), the underlying model’s reasoning capacity dictated the upper bound of success. Large frontier models consistently achieved perfect scores (6*/*6), whereas smaller, efficiency-focused models typically plateaued at 5*/*6, struggling with the most complex edge cases (Q6). The notable exception was Claude 3.5 Haiku, which matched the performance of significantly larger models, suggesting a highly efficient reasoning architecture suited for this specific agentic loop.

**Figure 3.**
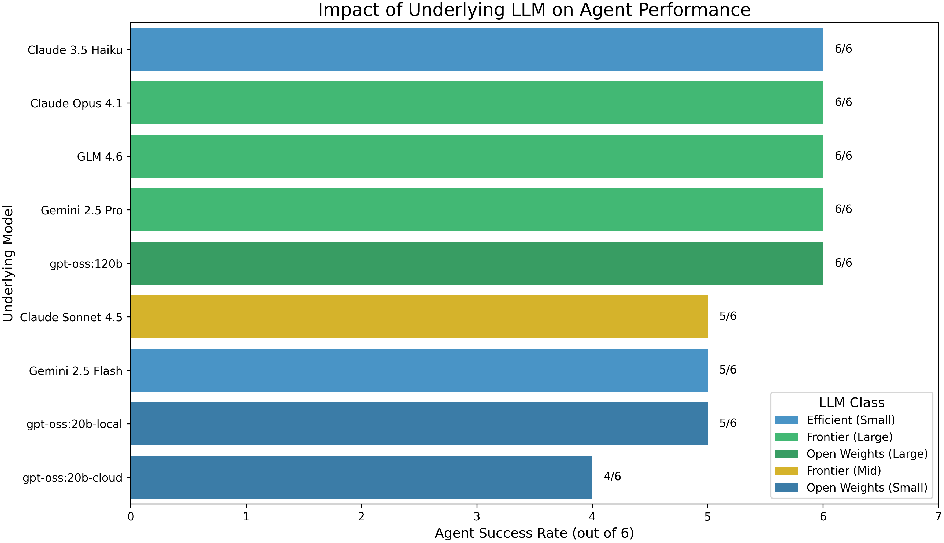
Performance of each LLM using GFFUtilsAI. This graphic demonstrates that while your agent framework raises the performance floor for everyone, the underlying LLM determines the ceiling.

### 5.3 Performance Impact of the Agentic Workflow

Upon integrating the models into our custom agentic workflow, we observed a dramatic and consistent performance improvement across all tested architectures. The agentic scaffolding effectively mitigated the reasoning failures observed in the zero-shot baseline. As detailed in Figure 4, the introduction of the agent resulted in a “Zero-to-Hero” effect for several models:GLM 4.6 and gpt-oss:120b, which both scored 0*/*6 in the baseline, achieved perfect scores across the 6-question test set within the agentic framework. Haiku 4.5 improved from 1*/*6 to 6*/*6, demonstrating that a lightweight model can achieve state-of-the-art accuracy when directed by a robust executive loop. The agentic workflow elevated the mean score of the test group from 0.8*/*6 (Baseline) to 5.6*/*6 (Agentic), representing a distinct architectural lift. The introduction of an agentic workflow transformed the performance landscape, essentially “fixing” the models that previously failed.

**Figure 4.**
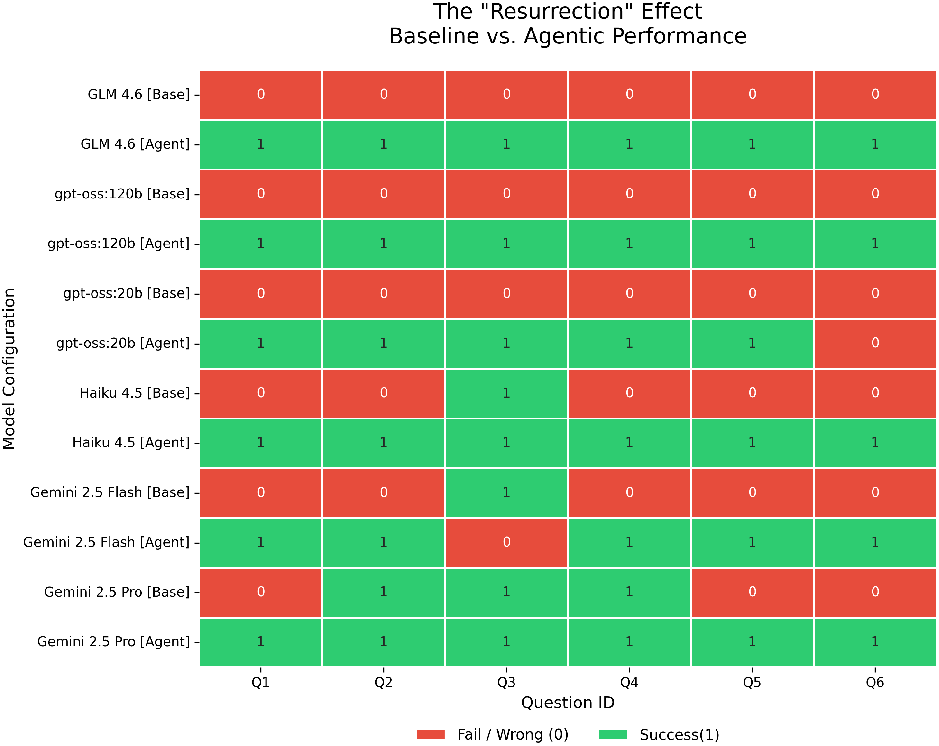
This heatmap shows the “Resurrection” Effect Baseline vs. Agentic Performance. This visual confirms that the failure mode of these models is likely executive rather than cognitive.

## 6 Discussion

GFFUtilsAI resolves complex queries that traditionally require significant time and coding expertise. While most standalone LLMs struggle to process GFF files directly, and the few capable competitors require up to eight minutes per query, GFFUtilsAI answers all test questions correctly in less than two minutes. Although currently designed for the specific task of exploring GFF files, this agentic architecture can be extended to broader bioinformatics domains, as demonstrated in recent studies [15, 16, 17].

The inherent stochasticity of LLMs poses a challenge to scientific reproducibility, influencing both tool selection and output formatting. While our findings indicate that agentic architectures offer greater stability than standalone models, the non-deterministic nature of these systems warrants caution. Consequently, users should validate findings through repeated execution to quantify potential variance before finalizing results.

A key avenue for future improvement is fine-tuning the underlying LLMs to enhance their ability to select the most appropriate tools for a given task.

We encourage interested researchers to assess the capabilities of GFFUtilsAI firsthand; the code is readily available for local deployment, and an interactive demonstration is accessible via a Jupyter Notebook environment hosted on the project’s GitHub repository.

## Declaration of AI Use

AI tools were utilized solely for English grammar refinement and as a coding copilot during software development. All scientific ideas, text generation, and final validation of results remain the exclusive responsibility of the authors.

## References

[1] Zoheir Ezziane. Applications of artificial intelligence in bioinformatics: A review. In Expert Systems with Applications 30 (2006) 2–10.

[2] Palistha Shrestha, Jeevan Kandel, Hilal Tayara and Kil To Chong. Post-translational modification prediction via prompt-based fine-tuning of a GPT-2 model. In Nature Communications volume 15, Article number: 6699 (2024).

[3] Manato Akiyama and Yasubumi Sakakibara. Informative RNA base embedding for RNA structural alignment and clustering by deep representation learning. In NAR Genomics and Bioinformatics, Volume 4, Issue 1, March 2022, lqac012.

[4] Zeming Lin, Halil Akin, Roshan Rao, Brian Hie, Zhongkai Zhu, Wenting Lu, Nikita Smetanin, Robert Verkuil, Ori Kabeli, Yaniv Shmueli, Allan dos Santos Costa, Maryam Fazel-Zarandi, Tom Sercu, Salvatore Candido and Alexander Rives. Evolutionary-scale prediction of atomic-level protein structure with a language model. In Science, 16 Mar 2023, Vol 379, Issue 6637, pp. 1123–1130.

[5] Kai Wang, Xuan Zeng, Jingwen Zhou, Fei Liu, Xiaoli Luan, Xinglong Wang. BERT-TFBS: a novel BERT-based model for predicting transcription factor binding sites by transfer learning. In Briefings in Bioinformatics, Volume 25, Issue 3, May 2024, bbae195.

[6] Haoran Chen, Shengxiao Zhang, Lizhong Zhang, Jie Geng, Jinqi Lu, Chuandong Hou, Peifeng He and Xuechun Lu. Multi role ChatGPT framework for transforming medical data analysis. In Scientific Reports volume 14, Article number: 13930 (2024).

[7] Taha ValizadehAslani, Yiwen Shi, Ping Ren, Jing Wang, Yi Zhang, Meng Hu, Liang Zhao, Hualou Liang. PharmBERT: a domain-specific BERT model for drug labels. In Briefings in Bioinformatics, Volume 24, Issue 4, July 2023, bbad226. 10.1093/bib/bbad226

[8] Isha Arora, Arkadij Kummer, Hao Zhou, Mihaela Gadjeva, Eric Ma, Gwo-Yu Chuang, Edison Ong. mtx-COBRA: Subcellular localization prediction for bacterial proteins. In Computers in Biology and Medicine, Volume 171, March 2024, 108114. 10.1016/j.compbiomed.2024.108114

[9] Jinhyuk Lee, Wonjin Yoon, Sungdong Kim, Donghyeon Kim, Sunkyu Kim, Chan Ho So and Jaewoo Kang BioBERT: a pre-trained biomedical language representation model for biomedical text mining. In Bioinformatics 36.4 (2020): 1234–1240.. 10.1093/bioinformatics/btz682

[10] Lin A, Ye J, Qi C, Zhu L, Mou W, Gan W, Zeng D, Tang B, Xiao M, Chu G, Peng S. Bridging artificial intelligence and biological sciences: a comprehensive review of large language models in bioinformatics. In Briefings in Bioinformatics. 2025 Jul;26(4):bbaf357. 10.1093/bib/bbaf357

[11] Dale R. gffutils. In https://github.com/daler/gffutils. Accessed 10-28-2025

[12] Patrick Gray, et al. strands-agents. In https://github.com/strands-agents/sdk-python Accessed: 12-2-2025.

[13] NCBI Resource Coordinators. Database resources of the National Center for Biotechnology Information Open Access In Nucleic Acids Research, Volume 41, Issue D1, 1 January 2013, Pages D8–D20.. 10.1093/nar/gks1189

[14] Peter J A Cock, Tiago Antao, Jeffrey T Chang, Brad A Chapman, Cymon J Cox, Andrew Dalke, Iddo Friedberg, Thomas Hamelryck, Frank Kauff, Bartek Wilczynski, Michiel J L de Hoon, Lin A, Ye J, Qi C, Zhu L, Mou W, Gan W, Zeng D, Tang B, Xiao M, Chu G, Peng S. Biopython: freely available Python tools for computational molecular biology and bioinformatics. In Bioinformatics. 2009 Mar 20;25(11):1422–1423.. 10.1093/bioinformatics/btp163

[15] Juexiao Zhou, Jindong Jiang, Zhongyi Han, Zijian Wang, Xin Gao. Streamline automated biomedical discoveries with agentic bioinformatics. In Briefings in Bioinformatics, Volume 26, Issue 5, September 2025.. 10.1093/bib/bbaf505

[16] Yang Liu, Rongbo Shen, Lu Zhou, Qingyu Xiao, Jiao Yuan, Yixue Li. A data-intelligence-intensive bioinformatics copilot system for large-scale omics research and scientific insights. In Briefings in Bioinformatics, Volume 26, Issue 4, July 2025, bbaf312.. 10.1093/bib/bbaf312

[17] K. Swanson, W. Wu, N.L. Bulaong et al. The Virtual Lab of AI agents designs new SARS-CoV-2 nanobodies. In Nature 646, 716–723 (2025).. 10.1038/s41586-025-09442-9

